# Fine-tuning STEAP1 protein expression and purification to preserve its conformation and function

**DOI:** 10.64898/2026.02.16.706263

**Authors:** Li He, Sungsoo Yoo, Hong Sun, Vasudha Pathakota, Mehma Kaur, Peng Li, Benjamin Alba, Xiaojie Yao

## Abstract

Six-transmembrane Epithelial Antigen of the Prostate 1 (STEAP1) has emerged as a promising therapeutic target for prostate cancer. We have optimized the expression and purification conditions of human STEAP1 to maximize the production of its homotrimeric form, which is crucial for metal ion reduction and maintaining cellular redox balance. Proteins obtained from these optimized conditions were complexed with both heme and flavin-adenine dinucleotide (FAD), two cofactors that are fundamental to STEAP functionality, suggesting native folding and interactions of the protein. In addition, we compared the impact of stable and transient expression systems on the protein quality of STEAP1. We found that stable expression promoted heme incorporation, improved expression homogeneity, and ensured correct protein orientation on cell surfaces. Our findings present effective strategies for optimizing the recombinant production of STEAP1, with potential applicability to other STEAP family proteins to facilitate therapeutic discovery.

## Introduction

Six-transmembrane Epithelial Antigen of the Prostate 1 (STEAP1) is a member of the STEAP protein family and plays a significant role in cellular processes, particularly in metal ion metabolism and redox reactions[1-3]. Initially identified for its overexpression in prostate cancer [4], STEAP1 has been the target of several therapeutic strategies, including antibody-drug conjugates (ADC), T cell engaging bispecific antibodies (T-BsAb), and chimeric antigen receptor (CAR) T cell therapies[5-8]. While there is limited STEAP1 expression in normal tissue, STEAP1 is overexpressed across various solid tumors, most notably in prostate cancer and Ewing sarcoma, indicating its broader role in tumorigenesis and cancer progression and making STEAP1 a highly promising therapeutic target for cancer treatment[9-13]. Moreover, STEAP1’s differential expression in cancerous and normal tissues makes it an important biomarker for cancer prognosis and diagnosis, as well as for monitoring disease progression[14-16].

The precise function of STEAP1 *in vivo* remains unclear. However, for other members of the STEAP family, namely STEAP2-4, in vivo metalloreductase activities have been measured[2, 17]. These activities involve the reduction of iron(III) and copper(II) ions via an electron transfer cascade, which requires three cofactors: NADPH, FAD and heme. NADPH binds to the cytosolic N-terminus oxidoreductase domain (OxRD) of STEAP, providing the necessary electron for this process[18, 19]. The electron is then transferred to FAD located at the cytosolic side of the transmembrane domain, from which the electron is next transferred to heme at the extracellular side of the transmembrane domain[20]. From heme, the electron may finally be transferred to a target substrate which will be reduced[21]. CryoEM structures of STEAP2 and STEAP4 suggest that STEAPs form trimers. Within the trimer, electrons may be transferred *in trans* from a neighboring OxRD of a STEAP monomer to the remaining cascade of a neighboring monomer[19, 22].

STEAP1, unlike STEAP2-4, possesses a significantly shorter N-terminal cytosolic domain, which prevents it from interacting with NADPH due to the absence of the OxRD in its N-terminus. The shorter FAD binding sequence also lowers its affinity for FAD. Without NADPH association, STEAP1 is unable to initiate the electron transfer cascade[2]. However, it acquires the reductase activity when fused to the N-terminal NADPH-binding domain of STEAP4[23], thus making it plausible that STEAP1 may become a functional reductase by complexing with other OxRD-containing STEAPs such as STEAP2[20].

The ability to obtain physiologically relevant purified STEAP1 proteins in their native and functional conformations is essential for the drug discovery process, facilitating target validation, compound or antibody screening, hit validation, structural studies, and precise characterization of drug-target interactions. Therefore, optimizing the expression and purification conditions of human STEAP1 to enhance proper folding and heme incorporation is highly desired. In this study, through inclusion of additives in culture media and using a stable expression system, we tailored conditions specifically for the optimal production of STEAP1, significantly enhancing trimerization and heme incorporation. We believe that this strategy could be extended to other heme-containing membrane proteins, thereby broadening its utility in protein research and therapeutic development.

## Results

### Optimization of transient expression conditions

The heme group is essential for the function of STEAP proteins, serving as a cofactor in their enzymatic reactions. A major challenge of producing heme-containing proteins like STEAP1 is ensuring the incorporation of the heme group into the protein during its synthesis and folding. To address this challenge, we first optimized the expression conditions of STEAP1 by examining the influence of recombinant expression culture harvest time and heme media additives.

Constructs encoding human STEAP1 with a C-terminal FLAG tag (referred to as STEAP1-FLAG, as depicted in the upper scheme of Fig. 1A) were transiently transfected into Expi293 GnTI-cells. Following the addition of enhancers, the cells were cultured at 37°C for varying durations (48, 72, or 96 hours). As illustrated in Fig. 1B, cell viability peaked at 48 hours and began to decline with extended culture time, with a more rapid decrease observed in cells expressing STEAP1 compared to those expressing an empty vector. In contrast, the viable cell density (VCD) remained stable for STEAP1-expressing cells, despite a steady VCD increase in cells expressing empty vectors. The lowered cell viability and VCD of STEAP1-expressing cells in comparison to control cells indicate a level of cytotoxicity associated with STEAP1 expression (Fig. 1B, dotted lines).

**Figure 1.**
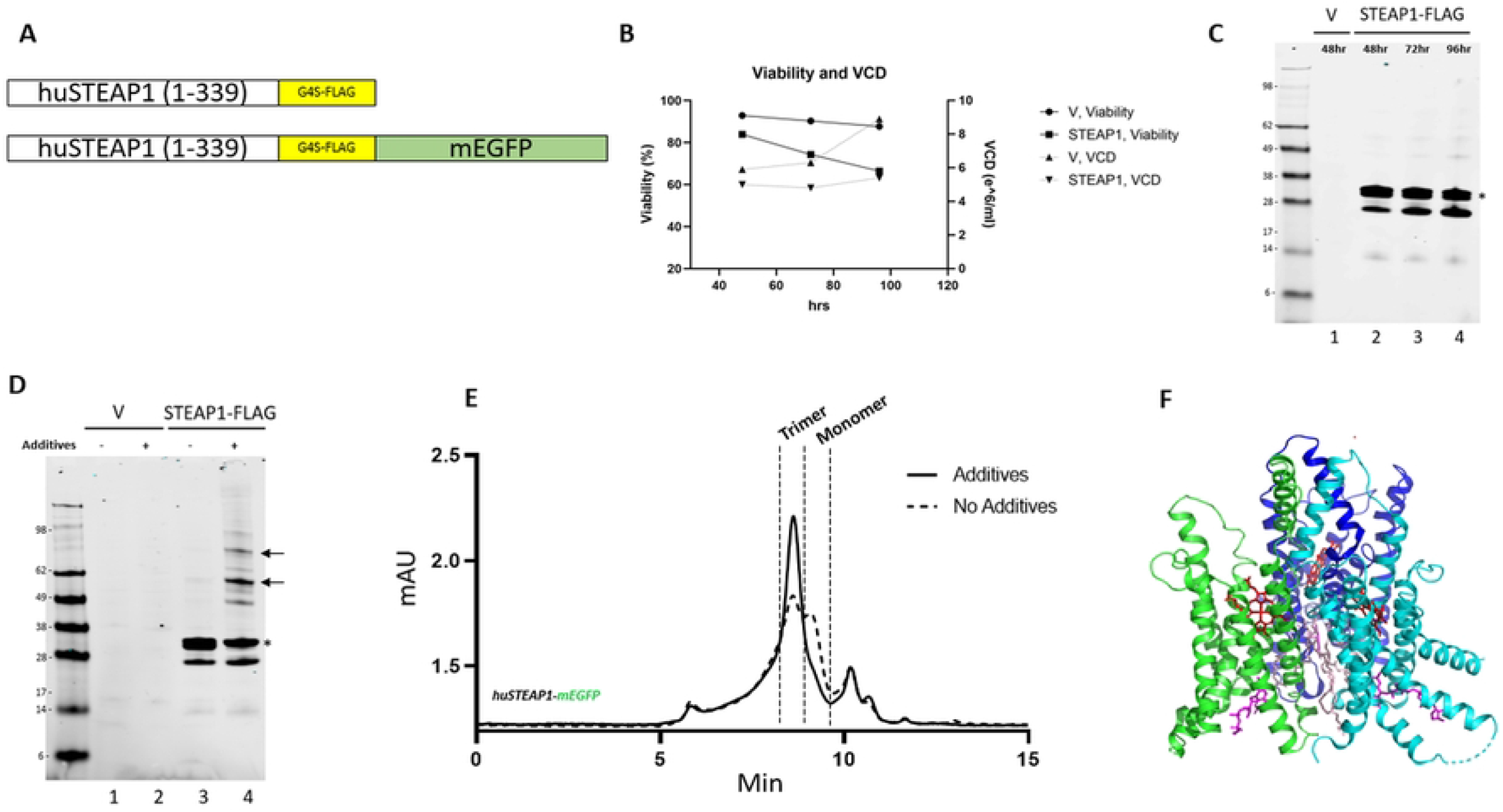
Optimization of transient expression conditions improved STEAP1 protein quality. A) Diagram of STEAP1 constructs. mEGFP: monomeric enhanced green fluorescent protein. B) Cell viability and VCD (Viable Cell Density) of cells expressing STEAP1 at different culturing timepoints. C) Western blot analysis of STEAP1 expression levels in relation to varying culturing durations. The asterisk denotes bands of STEAP1 positioned at the expected molecular weight. Arrows indicate bands of STEAP1 corresponding to the size of dimers and trimers. D) Measurement of STEAP1 expression levels in cells treated with heme additives (0.5 mM 5-Aminolevulinic acid HCl, 1 μM Hemin-Cl, and 5 μM Fe(III)Cl) or not via western blotting. An equal number of cells were used to prepare samples for each lane. E) FSEC analysis of the oligomeric status of STEAP1-mEGFP in cells treated with heme additives compared to those without additives. F) Cryo-EM structure of STEAP1 homotrimers (PDB: 8UCD), corresponding to the trimeric peak in Fig. 1E. The heme moiety is shown in red, lipid in pink, and bound FAD in magenta.

To determine whether prolonged culturing could compensate for the loss of cell viability through increased STEAP1 expression in live cells, we analyzed the expression levels of STEAP1 proteins via western blotting at various time points. Fig. 1C shows that samples from cells expressing STEAP1-FLAG consistently displayed similar levels of protein expression at the expected molecular weight (marked with “*”), indicating that extended culturing did not increase protein titers. Notably, a band of lower molecular weight became apparent with longer culturing times, suggesting that longer culturing might negatively impact protein homogeneity.

Previous reports suggest that supplementing compounds critical to heme biosynthesis boosted rabbit STEAP1 production in insect cells[20]. To test the effects of heme-supplemental factors in mammalian cells, we added 0.5 mM δ-aminolevulinic acid, 5 µM ferric chloride, and 1 µM hemin chloride (referred to as “additives”) to the cell culture 20 hrs post-transfection. Interestingly, the expression level of STEAP1, as measured by western blotting, was largely unaltered with or without additives (Fig. 1D, lanes #3 and #4). However, minor bands corresponding to the dimeric and trimeric forms of STEAP1 became more apparent when cells were supplemented with additives (Fig. 1D, lane #4). These oligomeric species resistant to denaturing SDS-PAGE conditions were also reported previously[20]. We further confirmed that the phenomenon was consistent, regardless of culturing time (data not shown). These results imply that while the additives did not significantly increase the total expression level of STEAP1, they may promote the formation of, or stabilize higher-order STEAP1 oligomers, which are crucial for the functional activity of STEAP1.

To verify whether the oligomer bands of STEAP1 observed in SDS-denaturing western blotting conditions are also present under non-denaturing conditions, Fluorescence-detection Size Exclusion Chromatography (FSEC) analyses were performed. An eGFP was fused to the C-terminus of STEAP1-FLAG (referred to as STEAP1-GFP, as depicted in the lower schematic of Fig. 1A). This fusion did not affect the expression level of STEAP1, as validated through western blot analysis (supplementary Fig. 1), and facilitated the monitoring of protein abundance via FSEC.

As shown in Fig. 1E, a clear bimodal distribution of STEAP1-GFP without heme additives (dotted black line) was observed, indicating the presence of two multimeric states. When heme additives were introduced (solid black line), there was a significant increase in the proportion of higher-order STEAP1 (first peak) relative to lower-order STEAP1 (second peak). Based on previous literature[20], we hypothesize that these two peaks represent trimeric and monomeric populations of STEAP1, respectively. Consistent with this hypothesis, human STEAP1 purified from the first peak was analyzed via cryoEM, yielding a high-resolution structure of trimeric STEAP1 (PDB: 8UCD) [8]. Importantly, both FAD and heme cofactors were clearly resolved within the transmembrane domain, indicating that the native cofactor binding and biological interactions of STEAP1 were preserved (Figure 1F)[8].

### Stably expressed STEAP1 shows increased heme incorporation

With the goal of optimizing STEAP1 protein production, we wanted to further optimize the culture conditions for STEAP1 expression. Stable cell pools are often preferred for large-scale protein production due to their consistent expression profiles. However, generating stable cell pools for membrane proteins can be challenging due to potential toxicity associated with their overexpression, which often leads to impaired cell growth.

Our first objective was to determine the feasibility of establishing a stable cell pool which expresses STEAP1. We co-transfected a STEAP1-FLAG construct along with a vector encoding PiggyBac transposase into Expi293 GnTI-cells. We then assessed whether stable expression of STEAP1 could achieve similar titer compared to transient transfection expression by western blotting. As shown in Fig. 2A, cells stably expressing STEAP1 exhibited a comparable, if not higher, expression level compared to those transiently expressing STEAP1 (lane #3 and #5) when lysates prepared from equal number of cells were loaded. Furthermore, when heme additives were provided during batch production, stably expressed STEAP1 mirrored the oligomer band profile observed with transiently expressed STEAP1, suggesting comparable protein integrity across both expression strategies.

**Figure 2.**
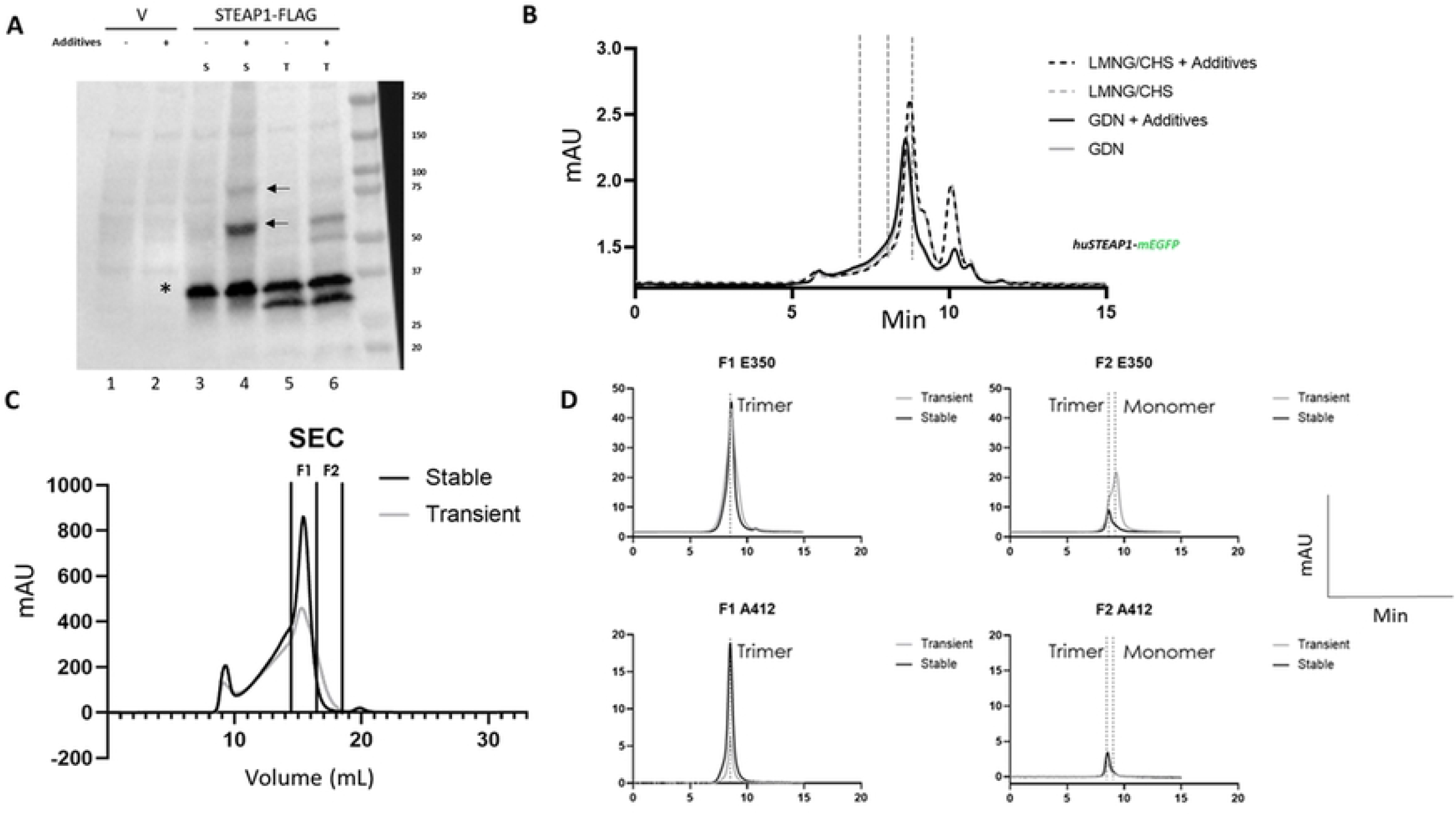
Enhancement of stable expression and detergent optimization significantly increased heme incorporation. A) Expression levels of STEAP1 in a stable expression system (S) and a transient expression system (T). An equal number of cells were used to prepare samples for each lane. Where indicated, heme additives were introduced during expression. The asterisk denotes bands of STEAP1 positioned at the expected molecular weight. Arrows indicate bands of STEAP1 corresponding to the size of dimers and trimers. B) Comparison of detergent conditions for STEAP1 solubilization through FSEC assays on STEAP1-mEGFP. Heme additives were added during solubilization where specified. C) SEC elution profiles of purified STEAP1 proteins from stable and transient expression systems. F1 represents pooled peak fractions indicative of the trimeric complex, while F2 denotes pooled fractions from the right shoulder of the peak. D) Measurement of heme incorporation into purified STEAP1 by emission at 350 nm and absorbance at 412 nm in SEC assays.

Before we compared the purified STEAP1 from stable versus transient expression systems, we first determined the optimal detergent condition for STEAP1 purification. Employing the FSEC system to monitor the quality of STEAP1 during solubilization, we compared two detergents that had been used for structural studies of STEAP1 proteins, namely LMNG/CHS[8] and digitonin[23] (Fig 2B), although we used GDN, a synthetic alternative to digitonin. As shown in Fig. 2B, using GDN for solubilization results in a more uniform oligomerization pattern by minimizing the hypothetical monomer peak. While the LMNG/CHS condition may produce a higher total yield as suggested by the larger area under the curve, this condition also increases heterogeneity with a more prominent monomer peak. Thus, we opted to use GDN as the detergent for purifying STEAP1. The effect of heme additives in the solubilization buffers was also tested. For each detergent condition, FSEC profiles were compared for STEAP1 solubilized with or without heme additives in the solubilization buffer. The presence of heme additives does not affect the quality of the solubilized STEAP1 proteins, as similar elution profiles were observed for both conditions (Fig. 2B). Thus, we decided to use GDN without additives as the solubilization buffer for the comparison study.

We then compared the quality of STEAP1 proteins purified from stable and transient expression systems. Fig. 2C illustrates the elution profile of affinity-purified STEAP1 proteins from analytical HPLC size exclusion chromatography (SEC). STEAP1 purified from a stable cell pool displayed a sharper elution profile compared to STEAP1 purified from the transient expression system. To further analyze the purified STEAP1 proteins, we pooled the peak fractions from each run, referred to as “F1”, and the right shoulder to the peak fraction, referred to as “F2”. F1s from each elution were subjected to HPLC SEC analyses. The absorbance at 412nm was monitored to assess the amount of heme incorporated into the purified STEAP1. Interestingly, we found several-fold higher heme incorporations in the F1 fraction of STEAP1 purified from stable cell pool compared to that from the transient expression system (Fig 2D bottom left, “F1 A412”). In contrast, the protein level, as measured by the emission spectra of tryptophan at 350nm, is comparable for the two F1 pools (Fig 2D top left, “F1 E350”). These results suggest that STEAP1 purified from a stable cell pool is significantly more efficient in heme incorporation.

We then subjected F2 pools to HPLC SEC analyses. Interestingly, a distinct peak matching the monomeric form of STEAP1 was observed only in the F2 fractions from the transient expression system. This contrasts with the elution profile of the F2 fractions from the stable expression system, where peak corresponding to the monomeric form of STEAP1 was barely visible (Fig 2D right top panel, “F2 E350”). Importantly, the peak attributed to the monomeric form of STEAP1 did not show heme absorbance, whereas only the trimeric form displayed a corresponding heme absorbance peak (Fig 2D right bottom panel, “F2 A412”). Thus, STEAP1 purified from the stable expression system resulted in a higher proportion of trimeric STEAP1, which, in turn, promoted heme incorporation since heme is incorporated only in the trimeric form.

### STEAP1 exhibits a more heterogeneous expression pattern in the transient expression system

Due to the observed variations in heme incorporation and protein behavior in SEC between transiently and stably expressed STEAP1, we utilized FACS to analyze the expression profile of STEAP1 across different expression systems. Cells were fixed, permeabilized, and treated with FITC-conjugated anti-FLAG antibodies 48 hrs post-transfection. As shown in Fig. 3, control cells without transfection exhibited a uniform peak with low mean fluorescence intensity (MFI) (∼5×10^2^) (Fig. 3A, orange). Stably expressing STEAP1-FLAG led to a ∼20-fold increase in MFI, shifting the peak to ∼10^4^ (Fig. 3A, blue vs. orange).

**Figure 3.**
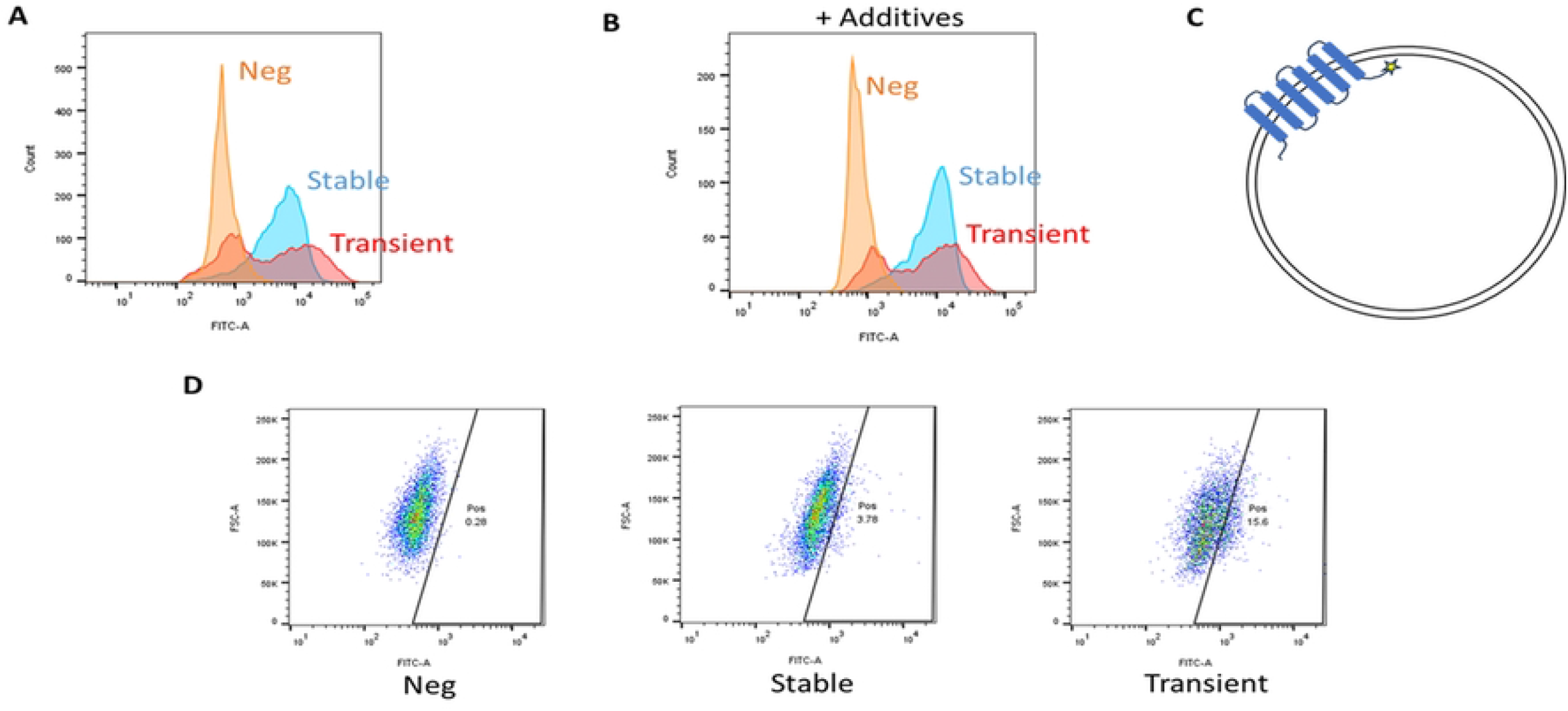
Flow cytometry analysis of STEAP1 expression in transient and stable expression systems. A) Flow cytometry analysis of total STEAP1 expression level in cells that were fixed and permeabilized. B) Same analysis as A), with the addition of heme additives during the expression. C) Orientation scheme of STEAP1 on the plasma membrane, noting that if correctly folded, the C-terminus of STEAP1 should face the cytoplasm and be inaccessible to antibodies. Star: C-terminus of STEAP1 protein. D) FACS analysis of FLAG epitope at the C-terminus exposed on the cell surface. Results are representative of two independent replicates. Neg: negative control cells with no STEAP1 expression. Stable: cells stably express STEAP1. Transient: cells stably express STEAP1.

Conversely, cells transiently expressing STEAP1 displayed a bimodal distribution of fluorescence signal, with approximately 30% of the cell population showing low MFI and the remaining ∼70% exhibiting high MFI, suggesting a ∼70% transfection efficiency. Notably, we constantly observed an ultra-high-MFI shoulder unique to transient transfection (Fig. 3A, red vs. blue), with MFI in this shoulder reaching up to ∼10^5^. Taken together, these data revealed a highly heterogeneous profile of expression among transient expression cell populations. Furthermore, they suggested that a substantial proportion of cells (∼15%) in the transient expression system exhibited protein production exceeding even the highest levels observed in stable cell pools.

We then investigated whether the inclusion of additives would impact the expression pattern of STEAP1. Cells transiently and stably expressing STEAP1 were cultured with heme supplementation for 48 hours, after which the total STEAP1 proteins in cells were measured by FACS. As shown in Fig. 3B, the expression pattern of both transient and stable systems remains unchanged. The results suggest that the addition of heme does not change the heterogeneous expression pattern observed among transient expression cells, although it promotes STEAP1 trimerization (Fig. 1E).

To conclude, we assessed the integrity of cellular STEAP1 expressions by staining the FLAG tags present on the cell surface in live cells. Under normal circumstances, the C-terminus of STEAP1 is in the cytoplasm, thus preventing access to the C-terminal FLAG tag for cell surface staining (Fig. 3C). However, when the cellular folding and quality control mechanisms are overwhelmed, misfolded or incorrectly oriented STEAP1 molecules may reach the cell surface, exposing the FLAG tag to the extracellular environment. As demonstrated in Fig. 3D, live cells were stained with FITC-conjugated anti-FLAG antibodies, followed by FACS analysis to detect exposed FLAG sequences. A more heterogeneous pattern was observed for transient expression cells compared to stable expression cells. By employing a gating named “Pos” to exclude nearly all cells in the negative sample, it was determined that approximately 3% of stable expression cells and 15% of transient expression cells could be stained.

## Discussion

The oligomerization of STEAP proteins is an important aspect of their functional regulation and activity[22]. Our study on human STEAP1 aligns well with previous studies[20] by highlighting the impact of cellular environment and expression conditions in preserving its native conformation. Importantly, these conditions also ensure the protein’s capacity to correctly incorporate heme, as observed in the intact heme and FAD cofactors in the resolved cryo-EM structure.

We also investigated how transient and stable expression systems impact trimer formation and co-factor incorporation of STEAP1. Our study indicates that transient expression systems may produce STEAP1 proteins in a shorter timeframe, but they may also result in incorrect orientation and less efficient heme incorporation in purified proteins. Conversely, stable expression systems, which integrate the gene-of-interest into the host genome, tend to produce protein at moderate and consistent levels, potentially facilitating proper membrane integration and heme incorporation.

Differences in protein production between transient and stable expression systems have been studied across various models[24, 25]. Although cells transfected either transiently or stably are expected to retain all necessary machinery for protein synthesis and quality control, the limited timeframe of transient expression may pose challenges for cells in adapting effectively to protein overexpression. This challenge might be more prominent for STEAP1 production, which requires coordination of heme incorporation, trimerization, and FAD association. We reasoned that these interdependent processes introduced additional complexity to STEAP1 complex formation, which transient expression systems may sometimes struggle to fully accommodate. The findings may offer valuable insights into the selection of expression systems for other similar proteins.

In conclusion, given the critical dependence of STEAP’s function on correct heme incorporation and protein conformation, the choice of expression system requires careful consideration in research and therapeutic discovery. It is also important to consider that, given the lengthy process and the potential toxicity to cells associated with membrane protein overexpression in stable cell line generation, transient expression remains the primary method in membrane protein research. This report demonstrates that, through meticulous optimization of both expression and purification conditions, well-folded and functional STEAP1 protein can be obtained. These results facilitate investigations into the structural-functional relationships underlying the metalloreductase activity of the STEAP1 protein, thereby enhancing the efficiency of therapeutic discovery.

## Materials and Methods

### STEAP1 construct design

All STEAP1 constructs were cloned into a proprietary expression vector compatible with a piggyBac transposase system (PB). DNA fragments encoding full-length human STEAP1 (1-339) followed by a G4S linker and a FLAG tag were synthesized by either IDT or Twist Bioscience and inserted into the expression vector using a Golden Gate Assembly method. When noted, a full eGFP sequence was fused to the C-terminal of the FLAG tag (FLAG-mEGFP). For stable cell transfection, a separate PB vector encoding the PB transposase was co-transfected with the expression vector containing the GOI. The expression vector also encodes a puromycin resistance gene. All DNA constructs underwent sequence verification prior to transfection.

### Cell culture and heme additives

Expi293 GnTI- cells were maintained in Expi293™ Expression Medium (cat. no. A1435102, Gibco). Cell viability and VCD were measured using Vi-cell BLU Cell Viability Analyzer (Beckman Coulter). The measurement was done using normal mode with 200 μL of sample.

Heme additives were added to cell culture when noted. The heme additives consisted of 0.5 mM 5-Aminolevulinic acid HCl (cat. no. A7793, Sigma), 1 μM Hemin-Cl (cat. no. 3741, EMD Millipore), and 5 μM Fe(III)Cl (cat. no. 710857, Sigma).

### STEAP1 transient and stable transfection

For transient transfection, Expi293 GnTI-cells were seeded at 3×10^6^ cells/mL with >90% viability at the day of transfection (day 0). Transfection was done using the ExpiFectamine™ 293 Transfection Kit (cat. no. A14525, Thermo Fisher Scientific) following the manufacturer’s protocol on day 0. Enhancers 1 and 2 were added to cells 20 hr post transfection. Cells were counted and harvested at 48 hrs, 72 hrs, or 96 hrs post transfection.

For stable transfection, Expi293 GnTI-cells were seeded at 0.75×10^6^ cells/mL density with >90% viability at the day of transfection (day 0). Cells were co-transfected with a PB vector and an expression vector encoding GOI using Lipofectamine LTX (cat. no. 15338–100, Gibco) following manufacturer’s protocol. For a 4 mL stable transfection, 2.5 μg of both GOI and PB vectors were added to 0.5 mL of Opti-MEM media (cat. no. 31985–070, Gibco) in a 24 well deep well plate (24-DWP). For 50% GOI transfection condition, 1.25 μg of GOI vector and 1.25 μg empty expression vector were used in place of 2.5 μg GOI vector. 10 μl Lipofectamine LTX in 0.5mL of Opti-MEM media was then added to mixed vectors. The entire mixture was incubated at room temperature (RT) for 15–20 min. 3×10^6^ viable cells were washed once with PBS, re-suspended in 1 mL of Opti-MEM, and were added to each well. Mixture of Lipofectamine LTX and vectors was added dropwise to the wells. Cells were shaken at 235 rpm for 5–6 h at 37°C with 5% CO_2_ before addition of 2 mL of growth media.

48 hrs post-transfection, cells were selected in selection media (growth media supplemented with 2 μg/mL puromycin (cat. no. A11138-03, Gibco)). Cell viability and VCD were measured every 2-3 days to determine the recovery progress. After selection, recovered cells were seeded at 1.0 – 1.5 ×10^6^ cells/mL in vented shake flasks for batch production. The same Expi293™ Expression Medium was used for production. Cells were shaken at 130 rpm and were harvested when the viability started to drop (usually at day 4-5).

### STEAP1 protein purification

To extract recombinant STEAP1-FLAG protein, cell pellets were hypotonically lysed in ice cold Lysis buffer (20 mM Tris pH 8.0, 1mM EDTA, Protease inhibitors (cat. no. A32963. Pierce™ Protease Inhibitor Tablets, EDTA-free, 1 tab/50 mL)), kept under gentle rotation for 15mins at 4°C. The lysed membranes were collected by spinning down at 14k rpm (23k x g) at 4°C for 10mins. The pelleted membranes were solubilized in solubilization buffer (20 mM Tris pH 8.0, 150 mM NaCl, 1% (w/v) lauryl maltose neopentyl glycol (LMNG) and 0.01% cholesterol hemisuccinate (CHS) or 20 mM Tris pH 8.0, 200 mM NaCl, 1% GDN), protease inhibitors (Pierce™ Protease Inhibitor Tablets, EDTA-free, 1 tab/50 mL)) and dounced on ice for 25 strokes then further solubilized for 2 hrs with rotation at 40k rpm at 4°C. The solubilized lysates were clarified by centrifugation 18k rpm (39k x g) for 30min at 4°C. STEAP1 was purified through batch binding with FLAG affinity beads and elution with 150 μg/mL FLAG peptide. STEAP1 was further purified by size-exclusion chromatography (SEC) using a mobile phase composed of 20 mM Tris pH 8.0, 200 mM NaCl and 0.01% GDN. Peak fractions were analyzed by HPLC (ex.280nm/em.350nm, 412nm Abs) and SDS-PAGE Coomassie staining.

### STEAP1-eGFP fluorescence-detection size-exclusion chromatography (FSEC)

Cell pellets transiently expressing STEAP1-eGFP were solubilized in either 1% LMNG/0.1% CHS detergent containing solubilization buffer (20mM Tris pH 8, 150mM NaCl, 1% LMNG, 0.1% CHS, protease inhibitors, with or without 10μM Hemin-Cl) or 1% GDN detergent containing solubilization buffer (20mM Tris pH 8, 150 mM NaCl, 1% GDN, protease inhibitors, with or without 10μM Hemin-Cl) for 2 hours rocking at 4°C, then cleared with ultracentrifugation. Sample from the cleared supernatant was injected onto SRT-C SEC-500 column (4.6×300 mm, Sepax) run at 0.35 mL/min with a mobile phase of 25 mM HEPES pH 7.5, 250 mM NaCl, and 0.005% GDN. eGFP signal was detected via fluorescence detector (ex480/em510; PMT = 14).

### SDS-page and western blotting analysis

The protein expression level of STEAP1 was determined via SDS-PAGE and western blotting techniques. In brief, around 20k cells per lane were pelleted and solubilized as specified in the Protein Purification and Quantification section. 4X Sample Buffer (cat. no. NP0007, Invitrogen) and 10X Sample Reducing Agent (cat. no. B0009, Invitrogen) were added to the solubilized proteins to achieve a 1X final concentration. Protein samples were then loaded onto a 4–12% Bis-Tris gel and electrophoresed in 1x MES running buffer (both from Invitrogen). A typical run was done at 160 Voltage for 50 min. Following SDS-PAGE, proteins were transferred to nitrocellulose membranes, which were blocked using Blocking Buffer for Fluorescent Western Blotting (cat. no. MB-070, Rockland Immunochemicals) and incubated with monoclonal anti-FLAG antibodies (cat. no. F1804, Sigma). After three washes in TBST (containing 0.05% (v/v) tween-20), the nitrocellulose membranes were probed with Alexa Fluor-680 conjugated secondary antibody (Thermo Fisher Scientific). The images of fluorescent western blot were acquired using an Odyssey® infrared imaging system (LI-COR Biosciences).

### STEAP1 detection by flow cytometry

Cells were harvested and washed two times in PBS through centrifugation and re-suspension. For cell surface staining, cells were gently re-suspended in a staining buffer (BD Bioscience) containing 1:500 of FITC-conjugated anti-FLAG antibody (cat. no. F4049, Sigma). The mixture was incubated for 30 min at 4°C in dark. Post-staining, cells were washed three times at 4°C prior to flow cytometric analysis. For intracellular staining (total staining), cells were washed, fixed, and permeabilized using Cell Fixation & Permeabilization Kit following manufacturer’s protocol (cat. no. ab185917, Abcam). After fixation and permeabilization, cells were stained as described above. Flow cytometric measurements were performed using a BD FACSymphony™ A1 Cell Analyzer (BD Biosciences), and the data were analyzed using FlowJo software (FlowJo, LLC).

## Supplementary Material

Figure S1 (File: Figure_S1.tif): Western blot analysis showing the expression levels of STEAP1 and STEAP1-mEGFP constructs discussed in 2.1.

## AUTHOR CONTRIBUTIONS

**L. He**: Investigation; visualization; formal analysis; writing – original draft; writing – review and editing. **S. Yoo**: Investigation; visualization; formal analysis; writing – original draft; writing – review and editing. **H. Sun**: Investigation; writing – original draft. **V. Pathakota**: Investigation. **M. Kaur**: Investigation. **P. Li**: Investigation. **B. Alba**: writing – review and editing. **X. Yao:** Conceptualization; writing – original draft; methodology; supervision; writing – review and editing

## ACKNOWLEDGMENTS

The authors would like to thank Nate Yoder, Patrick Cannon, Marisela Killian for technical support, and Fei Li for technical advice.

## Supplementary Material

**Supplementary Figure 1.**
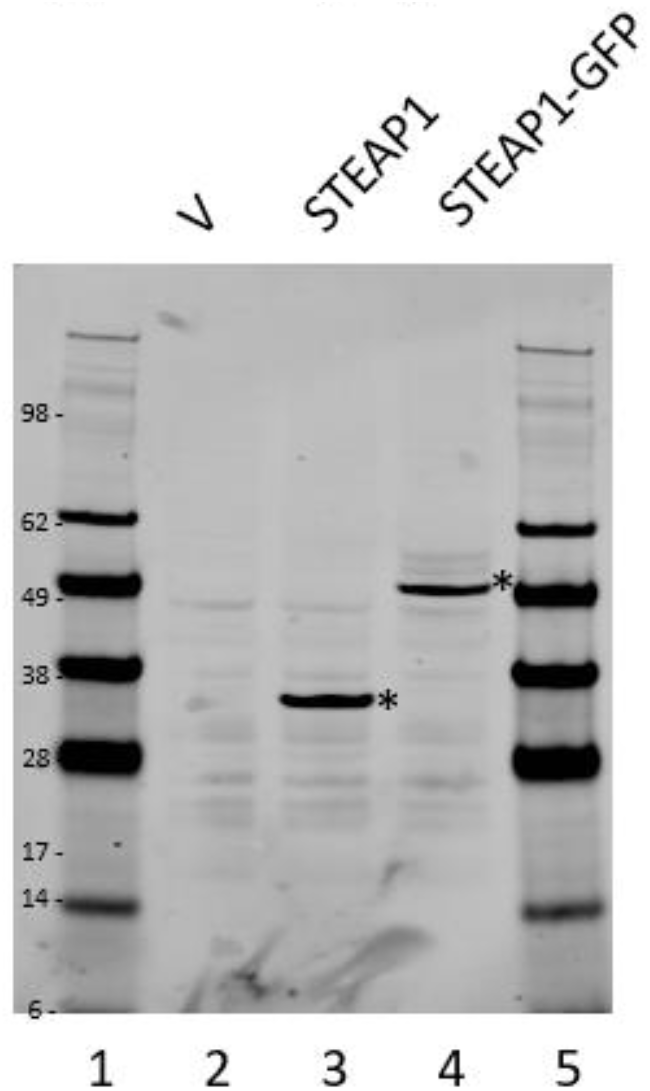
Western blot analysis showing the expression levels of STEAP1 and STEAP1-mEGFP constructs. The asterisk denotes bands of STEAP1 constructs positioned at the expected molecular weight.

## References

1. Grunewald, T.G., et al., The STEAP protein family: versatile oxidoreductases and targets for cancer immunotherapy with overlapping and distinct cellular functions. Biol Cell, 2012. 104(11): p. 641–57.

2. Ohgami, R.S., et al., The Steap proteins are metalloreductases. Blood, 2006. 108(4): p. 1388–94.

3. Cai, Q., et al., STEAP Proteins: Roles in disease biology and potential for therapeutic intervention. Int J Biol Macromol, 2025. 309(Pt 1): p. 142797.

4. Hubert, R.S., et al., STEAP: a prostate-specific cell-surface antigen highly expressed in human prostate tumors. Proc Natl Acad Sci U S A, 1999. 96(25): p. 14523–8.

5. Barroca-Ferreira, J., et al., Targeting STEAP1 protein in human cancer: current trends and future challenges. Current cancer drug targets, 2018. 18(3): p. 222–230.

6. Lin, T.Y., et al., Novel potent anti-STEAP1 bispecific antibody to redirect T cells for cancer immunotherapy. J Immunother Cancer, 2021. 9(9).

7. Nakamura, H., Y. Arihara, and K. Takada, Targeting STEAP1 as an anticancer strategy. Front Oncol, 2023. 13: p. 1285661.

8. Nolan-Stevaux, O., et al., AMG 509 (Xaluritamig), an Anti-STEAP1 XmAb 2+1 T-cell Redirecting Immune Therapy with Avidity-Dependent Activity against Prostate Cancer. Cancer Discov, 2024. 14(1): p. 90–103.

9. Moreaux, J., et al., STEAP1 is overexpressed in cancers: a promising therapeutic target. Biochemical and biophysical research communications, 2012. 429(3-4): p. 148–155.

10. Rocha, S.M., et al., The Usefulness of STEAP Proteins in Prostate Cancer Clinical Practice, in Prostate Cancer, S.R.J. Bott and K.L. Ng, Editors. 2021: Brisbane (AU).

11. Cheung, I.Y., et al., Novel markers of subclinical disease for Ewing family tumors from gene expression profiling. Clinical cancer research, 2007. 13(23): p. 6978–6983.

12. Fu, D., et al., A novel prognostic signature and therapy guidance for hepatocellular carcinoma based on STEAP family. BMC Med Genomics, 2024. 17(1): p. 16.

13. Burnell, S.E.A., et al., Utilisation of the STEAP protein family in a diagnostic setting may provide a more comprehensive prognosis of prostate cancer. PLoS One, 2019. 14(8): p. e0220456.

14. Wu, H.T., et al., The Tumor Suppressive Roles and Prognostic Values of STEAP Family Members in Breast Cancer. Biomed Res Int, 2020. 2020: p. 9578484.

15. Prasad, B., Y. Tian, and X. Li, Large-scale analysis reveals gene signature for survival prediction in primary glioblastoma. Molecular neurobiology, 2020. 57(12): p. 5235–5246.

16. Ohgami, R.S., et al., Identification of a ferrireductase required for efficient transferrin-dependent iron uptake in erythroid cells. Nature genetics, 2005. 37(11): p. 1264–1269.

17. Magnani, F., et al., Crystal structures and atomic model of NADPH oxidase. Proceedings of the National Academy of Sciences, 2017. 114(26): p. 6764–6769.

18. Chen, K., et al., Mechanism of stepwise electron transfer in six-transmembrane epithelial antigen of the prostate (STEAP) 1 and 2. Elife, 2023. 12.

19. Kim, K., et al., Six-transmembrane epithelial antigen of prostate 1 (STEAP1) has a single b heme and is capable of reducing metal ion complexes and oxygen. Biochemistry, 2016. 55(48): p. 6673–6684.

20. Kleven, M.D., M. Dlakic, and C.M. Lawrence, Characterization of a single b-type heme, FAD, and metal binding sites in the transmembrane domain of six-transmembrane epithelial antigen of the prostate (STEAP) family proteins. J Biol Chem, 2015. 290(37): p. 22558–69.

21. Oosterheert, W., et al., Cryo-EM structures of human STEAP4 reveal mechanism of iron (III) reduction. Nature communications, 2018. 9(1): p. 4337.

22. Oosterheert, W., et al., An Elegant Four-Helical Fold in NOX and STEAP Enzymes Facilitates Electron Transport across Biomembranes-Similar Vehicle, Different Destination. Acc Chem Res, 2020. 53(9): p. 1969–1980.

23. Baldi, L., et al., Recombinant protein production by large-scale transient gene expression in mammalian cells: state of the art and future perspectives. Biotechnology letters, 2007. 29(5): p. 677–684.

24. Chung, N.P., et al., Stable 293 T and CHO cell lines expressing cleaved, stable HIV-1 envelope glycoprotein trimers for structural and vaccine studies. Retrovirology, 2014. 11(1): p. 33.

